# SARS-CoV-2 Spreads through Cell-to-Cell Transmission

**DOI:** 10.1101/2021.06.01.446579

**Authors:** Cong Zeng, John P. Evans, Tiffany King, Yi-Min Zheng, Eugene M. Oltz, Sean P. J. Whelan, Linda Saif, Mark E. Peeples, Shan-Lu Liu

## Abstract

Severe acute respiratory syndrome coronavirus 2 (SARS-CoV-2) is a highly transmissible coronavirus responsible for the global COVID-19 pandemic. Herein we provide evidence that SARS-CoV-2 spreads through cell-cell contact in cultures, mediated by the spike glycoprotein. SARS-CoV-2 spike is more efficient in facilitating cell-to-cell transmission than SARS-CoV spike, which reflects, in part, their differential cell-cell fusion activity. Interestingly, treatment of cocultured cells with endosomal entry inhibitors impairs cell-to-cell transmission, implicating endosomal membrane fusion as an underlying mechanism. Compared with cell-free infection, cell-to-cell transmission of SARS-CoV-2 is refractory to inhibition by neutralizing antibody or convalescent sera of COVID-19 patients. While ACE2 enhances cell-to-cell transmission, we find that it is not absolutely required. Notably, despite differences in cell-free infectivity, the variants of concern (VOC) B.1.1.7 and B.1.351 have similar cell-to-cell transmission capability. Moreover, B.1.351 is more resistant to neutralization by vaccinee sera in cell-free infection, whereas B.1.1.7 is more resistant to inhibition by vaccine sera in cell-to-cell transmission. Overall, our study reveals critical features of SARS-CoV-2 spike-mediated cell-to-cell transmission, with important implications for a better understanding of SARS-CoV-2 spread and pathogenesis.

## INTRODUCTION

SARS-CoV-2 is a novel beta-coronavirus that is closely related to two other pathogenic human coronaviruses, SARS-CoV and MERS-CoV (Chan et al., 2020). The spike (S) proteins of SARS-CoV-2 and SARS-CoV mediate entry into target cells, and both use angiotensin-converting enzyme 2 (ACE2) as the primary receptor (Huang et al., 2020; Lan et al., 2020; Li, 2016; Walls et al., 2020; Zhou et al., 2020b). The spike protein of SARS-CoV-2 is also responsible for induction of neutralizing antibodies, thus playing a critical role in host immunity to viral infection (Barnes et al., 2020; Baum et al., 2020; Rogers et al., 2020; Zost et al., 2020).

Similar to HIV and other class I viral fusion proteins, SARS-CoV-2 spike is synthesized as a precursor that is subsequently cleaved and highly glycosylated; these properties are critical for regulating viral fusion activation, native spike structure and evasion of host immunity (Duan et al., 2020; Stewart-Jones et al., 2016; Sun et al., 2020; Watanabe et al., 2020; White et al., 2008). However, distinct from SARS-CoV, yet similar to MERS-CoV, the spike protein of SARS-CoV-2 is cleaved by furin into S1 and S2 subunits during the maturation process in producer cells (Chu et al., 2021; Coutard et al., 2020; Walls et al., 2020). S1 is responsible for binding to the ACE2 receptor, whereas S2 mediates viral membrane fusion (Shang et al., 2020; Wang et al., 2020). SARS-CoV-2 spike can also be cleaved by additional host proteases, including transmembrane serine protease 2 (TMPRSS2) on the plasma membrane and several cathepsins in the endosome, which facilitate viral membrane fusion and entry into host cells (Brooke and Prischi, 2020; Hoffmann et al., 2020; Lukassen et al., 2020).

Enveloped viruses spread in cultured cells and tissues via two routes: by cell-free particles and through cell-cell contact (Dale et al., 2011; Law et al., 2016; Mothes et al., 2010; Sattentau, 2008). The latter mode of viral transmission normally involves tight cell-cell contacts, sometimes forming virological synapses, where local viral particle density increases (Zhong et al., 2013b), resulting in efficient transfer of virus to neighboring cells (Mothes et al., 2010). Additionally, cell-to-cell transmission has the ability to evade antibody neutralization, accounting for efficient virus spread and pathogenesis, as has been shown for HIV and HCV (Brimacombe et al., 2011; Dale et al., 2013; Li et al., 2017; Miao et al., 2016; Zhong et al., 2013a). Low levels of neutralizing antibodies, as well as a deficiency in type I IFNs, have been reported for SARS-CoV-2 (Acharya et al., 2020; Jeyanathan et al., 2020; Lowery et al., 2021; Shang et al., 2020; Zhang et al., 2020b; Zhou et al., 2020a), and may have contributed to the COVID-19 pandemic and disease progression (Carvalho et al., 2021; Chu et al., 2020; Dispinseri et al., 2021; Hui et al., 2020; Park and Iwasaki, 2020; Yang et al., 2020).

In this work, we evaluated cell-to-cell transmission of SARS-CoV-2 in the context of cell-free infection and in comparison to SARS-CoV. Results from this in vitro study reveal the heretofore unrecognized role of cell-to-cell transmission that potentially impacts SARS-CoV-2 spread, pathogenesis and shielding from antibodies in vivo.

## RESULTS

### The spike protein of SARS-CoV-2 efficiently mediates cell-to-cell transmission of lentiviral pseudotypes

The spike is the only viral transmembrane protein that directly mediates SARS-CoV-2 entry into host cells. We evaluated if the spike protein of SARS-CoV-2 is critical for viral spread through cell-cell contact. In order to compare the efficiency of cell-to-cell vs. cell-free infection mediated by the spike proteins of SARS-CoV-2 and SARS-CoV, we took advantage of an intron-gaussia luciferase (inGluc) HIV-1 lentiviral vector bearing the spike of interest. In this system, the cells producing the inGluc lentiviral virions bearing the spike protein cannot themselves express Gluc because the intron is only removed during splicing of the virion genome transcribed from the integrated genome and not during the production of Gluc mRNA. However, when that lentivirus pseudotype enters a target cell, that genome is reverse transcribed and integrated in a new cell, and the CMV promotor drives transcription of the now intron-less Gluc transcript leading to Gluc protein production (Agosto et al., 2014; Mazurov et al., 2010). We measured Gluc activity as a readout to compare the cell-to-cell and cell-free infection efficiencies (Figure 1A **and** Figure 1B; see Methods). Because cell-contact-mediated infection comprises both cell-to-cell transmission and cell-free infection, we calculated the efficiency of cell-to-cell transmission by subtracting the portion of cell-free infection performed in parallel (see Methods).

**Figure 1.**
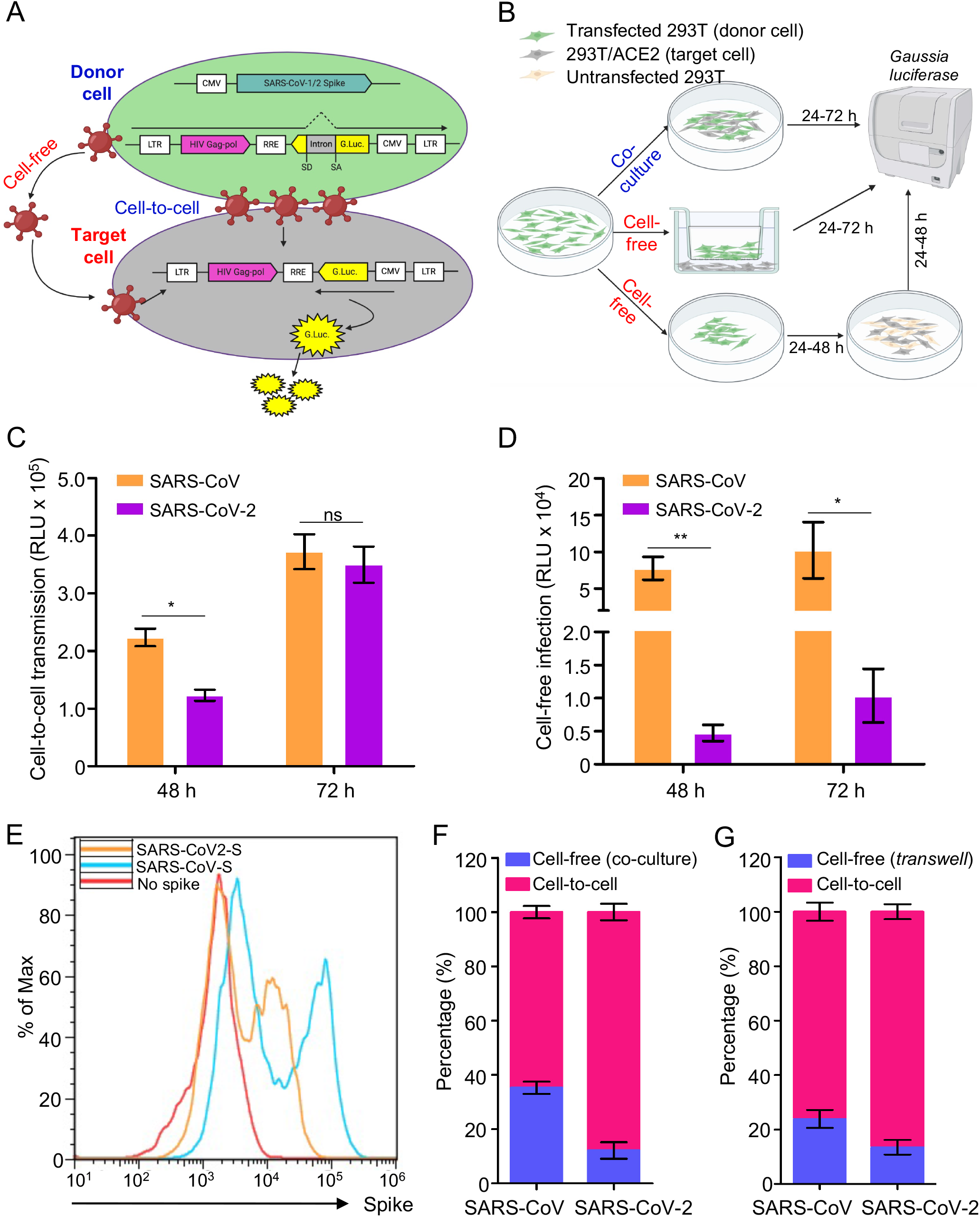
The spike protein of SARS-CoV-2 and SARS-CoV mediates cell-to-cell transmission of HIV-1 lentiviral pseudotypes. **(A and B)** Schematic representations of cell-to-cell and cell-free infection assays (see details in Methods). Briefly, the inGluc-based lentiviral pseudotypes bearing spike were produced in 293T cells, which were cocultured with the target cells (293T/ACE2) for cell-to-cell transmission; the Gluc activity of cocultured cells was measured over time (**A**). Cell-free infection was performed by harvesting virus from the same number of producer cells, followed by infecting 293T/ACE2 target cells in the presence of the same number of untransfected 293T cells; alternatively, cell-free infection was carried out in *transwell* plates, from which Gluc activity was measured (**B**). **(C)** Comparison of cell-to-cell transmission mediated by SARS-CoV-2 or SARS-CoV spike. Results shown were from 6 independent experiments, with cell-free infection measured at 48 and 72 hr after coculture; the portion of cell-free infection was excluded (n=6). **(D)** Comparison of cell-free infection mediated by SARS-CoV-2 or SARS-CoV spike. Results were from 6 independent experiments (n=6). **(E)** The expression level of spike proteins on the plasma membrane of donor cells was measured by flow cytometry using a polycolonal antibody T62, which detects both SARS-CoV-2 and SARS-CoV. **(F and G)** The calculated ratios between cell-to-cell and cell-free infection mediated by SARS-CoV-2 or SARS-CoV-2 spike. Results from cell coculture were shown in (F) and from *transwell* plates were shown in (G) (n=3∼6).

Despite a relatively low level of SARS-CoV-2 cell-to-cell transmission compared to SARS-CoV after 48 hr when coculturing of spike-bearing inGluc lentiviral pseudotype producer cells and 293T cells stably expressing human ACE2 (293T/ACE2), we observed similar levels of cell-to-cell transmission between SARS-CoV-2 and SARS-CoV by 72 hr, indicating a more efficient spread of SARS-CoV-2 (Figure 1C). In contrast, the rate of cell-free infection of SARS-CoV was much higher than that of SARS-CoV-2, i.e., approximately 10-fold, as measured at 48 and 72 hr post-infection (Figure 1D). Flow cytometric analysis of viral producer cells showed that the expression of SARS-CoV spike was higher than that of SARS-CoV-2 (Figure 1E), in agreement with our previous report (Zeng et al., 2020). By averaging results from six independent experiments, we estimated that cell-to-cell transmission contributed to >90% of the total SARS-CoV-2 spread in the coculturing system, as compared to ∼60% for SARS-CoV performed in identical experimental settings (Figure 1F). Parallel experiments were also performed by using a *Transwell* system, which showed ∼90% cell-to-cell vs. ∼10% cell-free infection for SARS-CoV-2 compared with ∼77% cell-to-cell vs. ∼23% cell-free for SARS-CoV (Figure 1G). Collectively, these results revealed that the spike protein of SARS-CoV-2 mediates cell-to-cell transmission more efficiently than the spike protein of SARS-CoV. However, the SARS-CoV spike is more capable of mediating cell-free infection compared with SARS-CoV-2.

### Recombinant VSV (rVSV) expressing SARS-CoV-2 spike spreads faster than rVSV bearing SARS-CoV spike

We next compared the spreading infection of replication-competent rVSV expressing SARS-CoV-2 or SARS-CoV spike. This system has been previously used to study the cell-cell transmission of Ebolavirus (EBOV) mediated by the glycoprotein, GP (Miao et al., 2016). Vero cells were inoculated with a relatively low MOI (0.01) of rVSV expressing GFP and SARS-CoV-2 spike in the place of VSV G protein (rVSV-GFP-SARS-CoV-2) or SARS-CoV spike (rVSV-GFP-SARS-CoV) (Case et al., 2020). Cells were overlayed by 1% methylcellulose to block viral diffusion, and the number and size of GFP-positive plaques were stained and determined by fluorescence microscopy. Despite similar numbers of GFP-positive plaques between SARS-CoV-2 and SARS-CoV, which confirmed equivalent inoculations, the sizes for SARS-CoV-2 plaques were noticeably larger, as inspected at 18 and 24 hr post-infection (Figure 2A **and** Figure 2B). Quantitative analyses of data at 72 hr showed that the size of SARS-CoV-2 plaques (diameter (0.93 ± 0.03) mm) was about 2 times greater than that of SARS-CoV (diameter (0.53 ± 0.02) mm), whereas the plaque numbers between SARS-CoV-2 and SARS-CoV were comparable (Figure 2C **and** Figure 2D).

**Figure 2.**
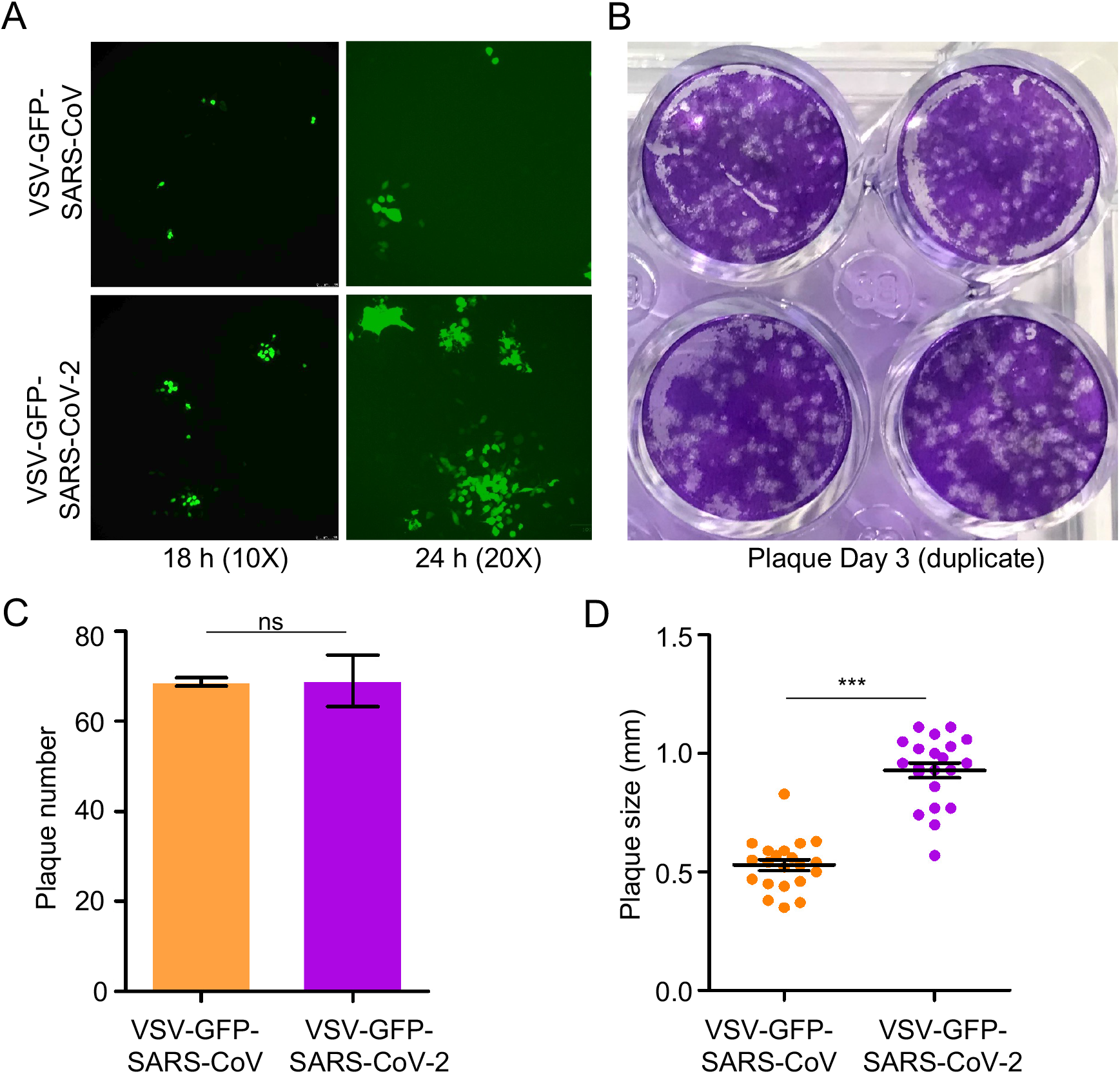
rVSV expressing SARS-CoV-2 spike spreads faster than does rVSV bearing SARS-CoV spike. Vero-E6 cells were infected with rVSV-GFP-SARS-CoV-2 or SARS-CoV (MOI=0.01); 1 h post-infection, cells were washed with PBS and cultured in the presence of 1% methylcellulose. Photos were taken at 18 h and 24 h **(A)**. After 72 hrs infection, cells were fixed with 3.7% PFA and stained with crystal violet **(B)**. The number and size of plaques were plotted in **(C) and (D),** respectively.

We next attempted to visualize cell-to-cell transmission of rVSV-GFP-SARS-CoV-2 by imaging fluorescent dye transfer in cocultured cells, either in the presence of methylcellulose or monoclonal antibody 2B04 against the SARS-CoV-2 spike. In this experiment, donor Vero cells were infected with rVSV-GFP-SARS-CoV-2 at different MOIs and subsequently cocultured with target Vero cells stably expressing mTomato (Vero-mTomato-Red). Efficient transmission was detected using fluorescence microscopy, as well as by flow cytometry at 6 h, with 23.9% double positive cell populations (Figure S1A **and** Figure S1B). Treating cocultured cells with methylcellulose, which has been found to prevent cell-free infection by drastically by reducing the diffusion of virions between cells (Mothes et al., 2010), or 2B04 that potently inhibit cell-free infection (Zeng et al., 2020), reduced the cell-to-cell transmission to 12.7% and 5.38%, respectively. Combining results from multiple independent experiments, we estimated that ∼50% of the total infection came from cell-to-cell transmission, which was still partially blocked by 2B04 (Figure S1C). Similar experiments performed in parallel for rVSV-GFP-SARS-CoV showed a stronger inhibition by methylcellulose (∼65%), suggesting a more efficient cell-free infection of rVSV-GFP-SARS-CoV compared with that of SARS-CoV-2. Importantly, 2B04 had no effect on cell-to-cell or cell-free infection of rVSV-GFP-SARS-CoV as would be expected since 2B04 does not cross-react with SARS-CoV (Figures S1D-S1F) (Alsoussi et al., 2020; Zeng et al., 2020). Altogether, these results demonstrated that, similar to lentiviral pseudotypes, the spike protein of SARS-CoV-2 more efficiently mediates the cell-to-cell transmission of rVSV-GFP than SARS-CoV.

### The higher cell-cell fusion activity of SARS-CoV-2 spike contributes to efficient cell-to-cell transmission

We next explored if cell-cell fusion by SARS-CoV-2 spike plays a role in cell-to-cell transmission. To this end, we co-transfected 293T cells with plasmids expressing the inGluc lentiviral vector, SARS-CoV-2 or SARS-CoV spike, and GFP. The transfected producer cells were cocultured with target 293T/ACE2 cells; syncytia formation and cell-to-cell transmission were measured over time. Following ∼2 h of coculturing, we observed small but apparent syncytia for SARS-CoV-2, yet with no syncytia formation for SARS-CoV (Figure 3A). At 24 h following coculturing, more syncytia formation, with larger sizes, was observed in cells expressing SARS-CoV-2 spike, whereas fewer and smaller syncytia were seen for SARS-CoV (Figure 3A). The difference between SARS-CoV-2 and SARS-CoV spike-induced cell-cell fusion was further evaluated by a more quantitative, Tet-off-based fusion assay, which showed a ∼5-fold higher fusion activity of SARS-CoV-2 compared with that of SARS-CoV (Figure 3B).

**Figure 3.**
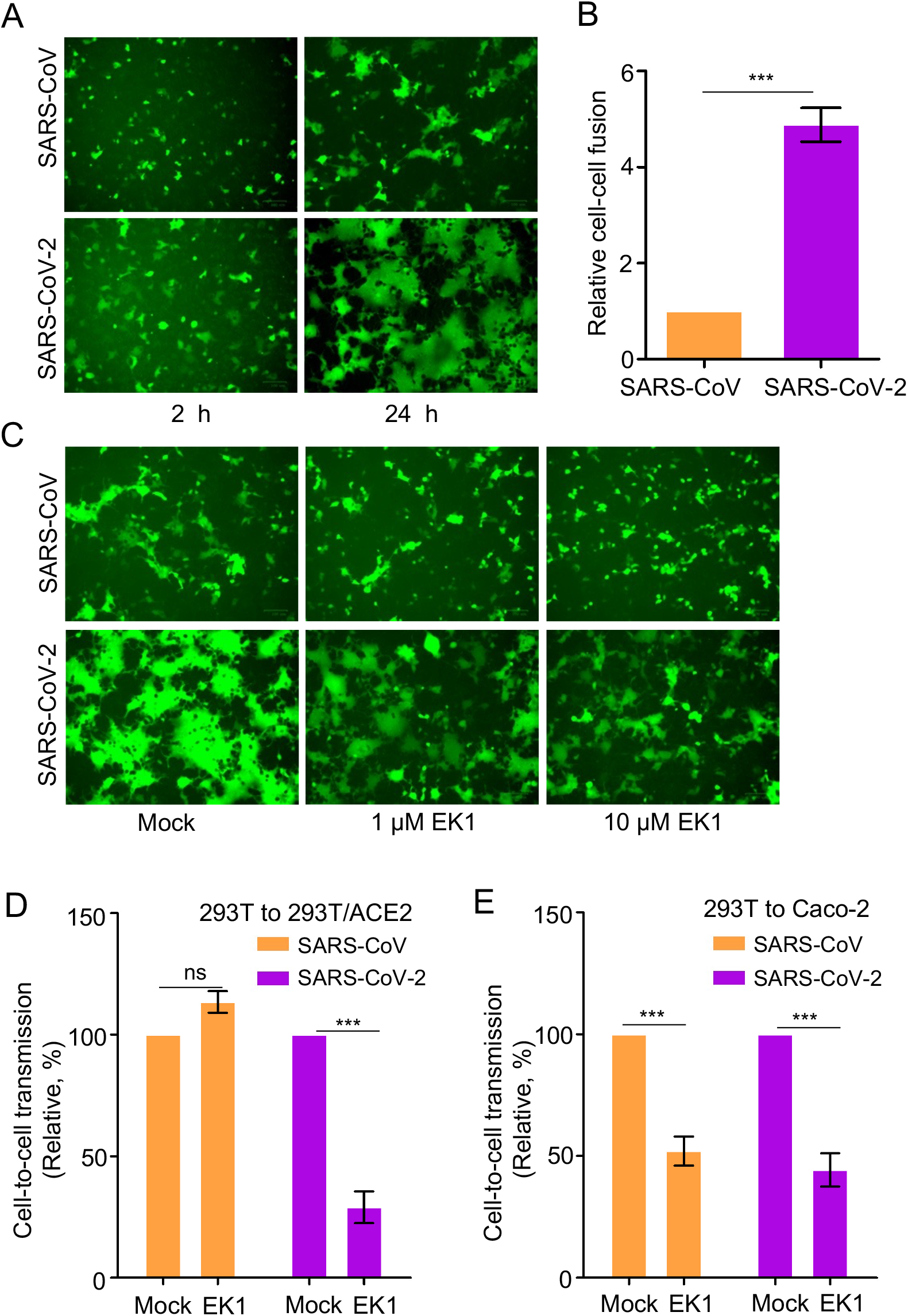
Cell-cell fusion mediated by SARS-CoV and SARS-CoV-2 spike contributes to cell-to-cell transmission. **(A)** Syncytia formation mediated by the spike of SARS-CoV-2 or SARS-CoV. 293T donor cells were cotransfected with plasmids encoding SARS-CoV-2 or SARS-CoV spike, lentiviral NL4-3 inGluc vector and eGFP. After 24 h post-transfection, the donor cells were cocultured with target 293T/ACE2 cells at 1:1 ratio, with fusion monitored over time and photos taken after 2 hr and 24 hr, respectively. **(B)** Quantification of cell-cell fusion. 293T cells were transfected with plasmids encoding tet-off or SARS-CoV or SARS-CoV-2 spike and were cocultured with target 293FT-mCAT-Gluc cells, which were transfected with a plasmid expressing ACE2; Gluc activity was measured from the supernatant of cocultured cells at 24 hr and 48 hr, respectively. Relative fusion was plotted by setting the fusion activity of SARS-CoV as 1.0. **(C, D and E)** Fusion inhibitor EK1 inhibits cell-cell fusion of SARS-CoV-2 spike, in accordance with its effect on cell-to-cell transmission. Effect of EK1 on syncytia formation induced by SARS-CoV-2 spike (**C**); photos were taken at 24 hr. Effects of EK1 on SARS-CoV-2 or SARS-CoV infection from 293T to 293T/ACE2 **(D)** or from 293T to Caco-2 **(E)**. Transfected 293T donor cells were cocultured with 293T/ACE2 or Caco-2 cells in the presence or absence of 10 µM EK1 and Gluc activity was measured at 24 to 72 hr after coculture. Results from 3 to 6 independent experiments were averaged and plotted as relative values by setting the mock control as 100% (n=3∼6).

We next treated cocultured cells with a pan-coronavirus fusion peptide inhibitor EK1 that has been shown to inhibit fusion of SARS-CoV-2, SARS-CoV, and other related CoVs (Xia et al., 2019; Xia et al., 2020), and simultaneously measured its effect on cell-cell fusion and cell-to-cell transmission. Syncytia formation of SARS-CoV-2 was strongly inhibited by EK1 (Figure 3C), in accordance with its effect on cell-to-cell transmission (Figure 3D). Unexpectedly, although EK1 inhibited the ability of SARS-CoV spike to induce small syncytia, we did not find obvious inhibition of EK1 on SARS-CoV spike-mediated cell-to-cell transmission (Figure 3C **and** Figure 3D). To investigate if these results were cell-type dependent, we performed similar experiments using human intestine epithelial Caco-2 as target cells and found that EK1 indeed inhibited the cell-to-cell transmission of both SARS-CoV-2 and SARS-CoV (Figure 3E). Overall, these results support the notion that the strong cell-cell fusion activity of SARS-CoV-2 spike contributes, but may not solely determine, its efficient cell-to-cell transmission.

### ACE2 enhances but is not required for cell-to-cell transmission

ACE2 is the primary receptor of both SARS-CoV-2 and SARS-CoV, mediating viral entry into host cells. We next evaluated the role of ACE2 in cell-to-cell transmission as compared with cell-free infection. We observed increased cell-to-cell and cell-free infection when more plasmid encoding ACE2 was transfected into the target 293T cells, as would be expected (Figure 4A and Figure 4B). Interestingly, with a relatively low dose of ACE2 (i.e., 0.2 μg), SARS-CoV-2 reached ∼70% of its maximal cell-to-cell transmission (at 0.5 μg ACE2). In contrast, SARS-CoV showed ∼30% maximal cell-to-cell transmission at 1.5 μg ACE2 (Figure 4A **and** Figure 4B). Notably, when the highest dose of ACE2 (1.5 μg) was transfected into target cells, we consistently observed decreased cell-to-cell transmission of SARS-CoV-2 compared with a continually increasing trend for SARS-CoV (Figure 4A **and** Figure 4B). This pattern of cell-to-cell transmission was different from that of cell-free infection, where both SARS-CoV-2 and SARS-CoV exhibited an increase, with similar kinetics, in a strictly ACE2 dose-dependent manner (Figure 4A **and** Figure 4B). We confirmed ACE2 expression in target cells by flow cytometry and western blotting (Figure S2A **and** Figure S2B). Consistent with increasing expression of ACE2 in target cells, we observed increasing sizes of syncytia formation for SARS-CoV-2, but cell-cell fusion by SARS-CoV was not evident (Figure S2C). Giant syncytia formation at 1.5 μg ACE2 resulted in cell death, which might have contributed to decreased cell-to-cell transmission for SARS-CoV-2 (Figure S2C). Overall, these results indicate that ACE2 enhances cell-to-cell transmission of both SARS-CoV-2 and SARS-CoV, yet the former requires less ACE2 for the process to occur.

**Figure 4.**
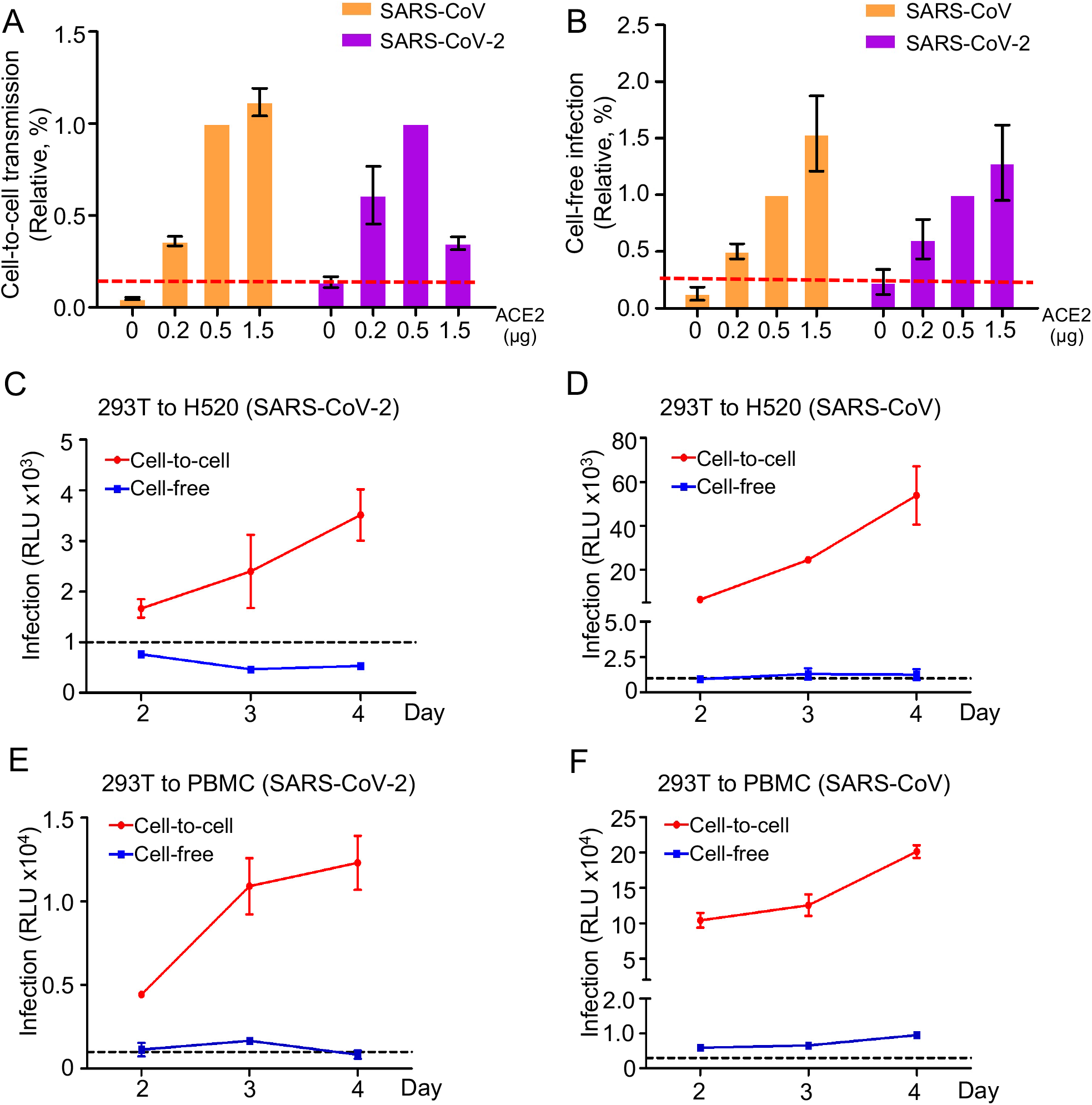
ACE2 enhances cell-to-cell transmission but is not absolutely required. (**A and B**) Cell-to-cell and cell-free infection was performed as described for Figures 1 and 3 except that target cells were 293T transfected with different amounts of a plasmid encoding ACE2. Relative rates of cell-to-cell transmission and cell-free infection were calculated by setting the values of 0.5 µg ACE2 to 1.0 **(A and B,** n=3). (**C, D, E and F**) Experiments were carried out as described for Figures 1 and 3 except that target cells were H520 and human PBMCs (n=3 for each).

We further explored if cell-to-cell transmission of SARS-CoV-2 can occur in the absence of ACE2 expression in target cells. We first used NCI-H520, a human lung epithelial cell line that expresses an extremely low level of ACE2 (Figure S2D). Cell-to-cell transmission was detected at day 2, which continued to increase through day 4. In contrast, cell-free infection was not detected in NCI-H520 cells throughout the 3-day period (Figure 4C). Cell-to-cell transmission of SARS-CoV was also observed in H520 cells, at a higher level than that of SARS-CoV-2; again, similar to SARS-CoV-2, no/low cell-free infection was detectable (Figure 4D). We next tested human PBMCs, which do not express ACE2 (Figure S2D), and observed apparent cell-to-cell transmission for both SARS-CoV and SARS-CoV-2, yet no/low cell-free infection was detected, the latter being consistent with recently published results (Banerjee et al., 2020) (Figure 4E **and** Figure 4F). As a control, we carried out cell-to-cell transmission and cell-free infection in Calu-3, a human lung epithelial cell line which expresses a high level of ACE2 (Figure S2D). A rapid increase in cell-to-cell transmission was observed for SARS-CoV-2 from day 2 through day 4, despite an overall level of infection for SARS-CoV that was higher than observed for SARS-CoV-2 (Figure S2E **and** Figure S2F). Together, these results demonstrated that cell-to-cell transmission of SARS-CoV-2 and SARS-CoV can occur in the absence of ACE2.

### Cell-to-cell transmission of SARS-CoV-2 involves endosomal entry

SARS-CoV-2 uses different pathways for entry, either at the plasma membrane and/or in the endosomal compartment (Harrison et al., 2020; Hoffmann et al., 2020; Murgolo et al., 2021; V’Kovski et al., 2021; Wrapp et al., 2020; Yeung et al., 2021). While our results indicated that entry via the plasma membrane is important for cell-to-cell transmission, we probed whether fusion in the endosomal compartment may also be involved. We applied in parallel a panel of endosomal inhibitors to the cell-to-cell and cell-free infection assays. We found that cathepsin L inhibitor III, cathepsin B inhibitor CA-074, E-64d (general cathepsin inhibitor), BafA1 (ATPase pump inhibitor), and Leupeptin (general protease inhibitor), all significantly inhibited cell-to-cell transmission (Figure 5A). Interestingly, the effect of these drugs on SARS-CoV-2 were generally less potent compared to SARS-CoV, with the exception of cathepsin L inhibitor III (Figure 5A). Moreover, these drugs generally showed a stronger effect on cell-free infection, again especially for SARS-CoV (Figure 5B). Of note, CA-074 had modest effects on both viruses (Figure 5B), which was consistent with the notion that cathepsin B does not play a significant role in cleaving the spike protein of SARS-CoV and SARS-CoV-2, which is required for fusion (Nitulescu et al., 2020; Ou et al., 2020). We also applied these inhibitors to cell-cell fusion assays but found no effect on either SARS-CoV-2 or SARS-CoV, as would be expected (Figure S3). To assess possible cell type-dependent effects, we carried out experiments using Caco-2 target cells and found that cathepsin L inhibitor III and BafA1 robustly inhibited cell-to-cell transmission and cell-free infection of both viruses, in particular SARS-CoV (Figure 5C **and** Figure 5D). Overall, these results support the notion that endosomal entry is involved in cell-to-cell transmission of SARS-CoV-2, and to a greater extent, SARS-CoV.

**Figure 5.**
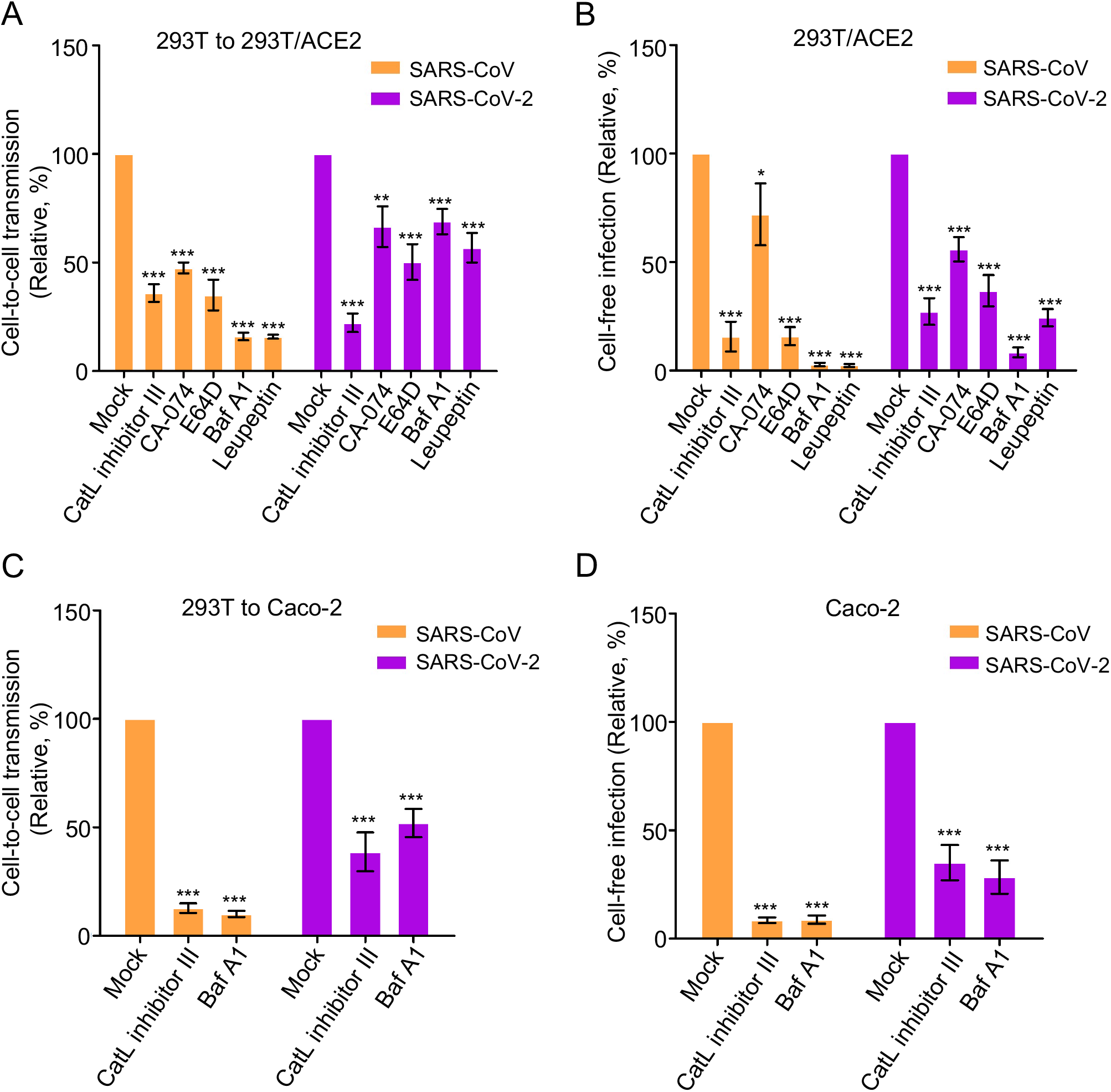
Endosomal entry pathway is involved in cell-to-cell transmission. Effect of endosomal entry inhibitors on cell-to-cell and cell-free infection of SARS-CoV-2 and SARS-CoV. Experiments were carried out as described in Figures 1C and 1D, except that indicated inhibitors were present during the infection period. The concentrations of inhibitors used were as follows: 1 µM or 5 µM Cat L inhibitor III, 1 µM or 5 µM CA-074, 10 µM or 30 µM E-64D, 25 nM or 50 nM BafA1, and 20 µM or 50 µM leupeptin. **(A and B**) Effect in 293T cells. **(C and D)** Effect in Caco-2 cells. In all experiments, Gluc activity was measured at 48 and 72 hr after infection, and rates of relative infection were plotted by setting the values of mock infection without drugs to 100. Results were from 4∼6 independent experiments.

### Cell-to-cell transmission of SARS-CoV-2 is refractory to neutralizing antibody and convalescent plasma

One important feature of the virus cell-to-cell transmission is evasion of host immunity, particularly neutralizing antibody-mediated response. We therefore examined the sensitivity of SARS-CoV-2 spike-mediated cell-to-cell transmission to neutralization by a monoclonal antibody against the receptor-binding domain of the spike, 2B04 (Alsoussi et al., 2020), as well as convalescent plasma derived from COVID-19 patients (Roback and Guarner, 2020; Zeng et al., 2020). While 2B04 effectively inhibited cell-free infection of SARS-CoV-2 in 293T/ACE2 cells by more than 90%, its effect on cell-to-cell transmission between 293T and 293T/ACE2 was ∼50% (Figure 6A **and** Figure 6B). As would be expected, 2B04 had no effect on SARS-CoV, regardless of cell-to-cell transmission or cell-free infection (Figure 6A **and** Figure 6B). We also performed cell-cell fusion in the presence of different concentrations of 2B04, and we found that the fusion activity of the SARS-CoV-2 spike was inhibited in a dose-dependent manner (Figure 6C). We then tested five serum samples of COVID-19 patients, and observed that, although they potently inhibited the cell-free infection of SARS-CoV-2 (p<0.001), they showed variable but no significant effect on cell-to-cell transmission of SARS-CoV-2; the effect of these sera on SARS-CoV infection, either cell-to-cell or cell-free, was minimal or modest (Figure 6D **and** 6E). Together, these results indicate that cell-to-cell transmission of SARS-CoV-2 is mostly refractory to neutralization by neutralizing antibodies against spike relative to cell-free infection.

**Figure 6.**
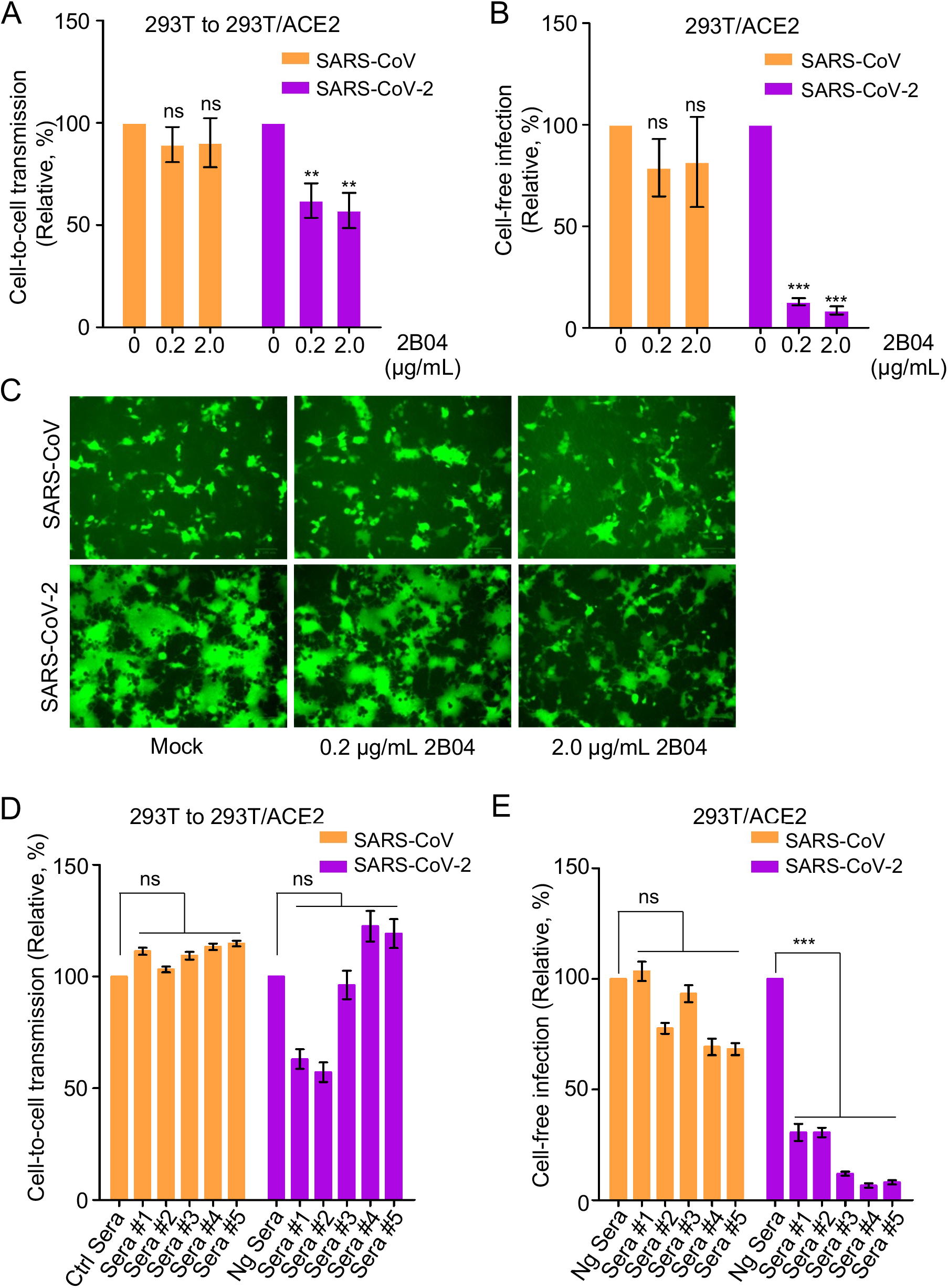
Cell-to-cell transmission of SARS-CoV-2 is refractory to inhibition by neutralizing antibody and COVID-19 convalescent plasma. **(A, B and C)** Effects of SARS-CoV-2 monoclonal antibody 2B04 on cell-to-cell transmission, cell-free infection, and cell-cell fusion mediated by SARS-CoV-2 or SARS-CoV-2 spike. The experiments were carried out as described in Figures 1C and 1D, except that 0.2 µg/mL or 2 µg/mL 2B04 were included during the infection period. Relative infections were plotted by setting the values of mock infection without 2B04 to 100% for statistical analyses (**A and B**). The photos of syncytia formation were taken at 18 h after coculture and presented (**C**). **(D and E)** Effect of COVID-19 sera on cell-to-cell and cell-free infection of SARS-CoV-2 and SARS-CoV. Experiments were performed as described as above, except five diluted COVID-19 sera were included during the infection period. Effect on cell-to-cell **(D)** and cell-free **(E)** of SARS-CoV or SARS-CoV-2 were summarized and plotted by setting the values of mock infection control to 100% (n=3∼4).

### Cell-to-cell transmission of SARS-CoV-2 variants of concern and their sensitivity to COVID-19 vaccinee sera

The D614G mutation in SARS-CoV-2 spike, as well as emerging variants of concern (VOCs) containing D614G and other key spike mutations, have been reported to enhance viral infectivity, transmissibility, and resistance to COVID-19 vaccines (Khan et al., 2021; Noh et al., 2021; Plante et al., 2021; Wu et al., 2021; Xie et al., 2021; Zhou et al., 2021). As such, we examined the cell-to-cell transmission capability of authentic SARS-CoV-2 WT (USA-WA1/2020), D614G variant (B.1.5), and two VOCs B.1.1.7 (501Y.V1) and B.1.351 (South African, 501Y.V2), in the presence or absence of pooled sera from mRNA vaccinees (3 Moderna and 3 Pfizer). Donor Vero-ACE2 cells were first infected with WT SARS-CoV-2 (MOI=0.2), D614G (MOI=0.02), B.1.1.7 (MOI=0.02), and B.1.351 (MOI=0.02), respectively. Note that a 10-fold higher MOI was used for WT in order to achieve comparable rates of infection in donor cells between WT and VOCs, given that D614G-containing variants are known to significantly increase the viral infectivity (Plante et al., 2021; Zhang et al., 2020a). Approximately 20 hrs post-infection, the culture media of donor cells was harvested, of which the whole volume of which was used to infect target Vero-mTomato-Red cells for 6 hr in order to determine the viral infectivity. In parallel, the infected donor Vero-ACE2 cells were digested, and cocultured with the same number of Vero-Tomato-Red cells as was used in the cell-free infectivity assay, also for 6 hrs, as a measurement of cell-to-cell transmission. To determine the sensitivity of cell-to-cell transmission vs. cell-free infection to neutralization by vaccinee sera, we pooled the serum samples of 6 mRNA vaccinees, i.e., 3 from Moderna and 3 from Pfizer, and added them to the cultured medium. The efficiency of cell-to-cell transmission and cell-free infectivity was determined by measuring the percentage of SARS-CoV-2 nucleocapsid (N)-positive Vero-mTomato-Red cells using flow cytometry. Considering the potential impact of infected donor cells on cell-to-cell transmission, we normalized the rate of cell-to-cell transmission with the total rate of virus spread in both SARS-CoV-2-positive Vero-mTomato-Red cells as well as Vero-ACE2 cells over the entire infection period, i.e., from the initial infection of donor cells to the end of coculture (∼26 hrs).

Representative flow cytometric results and summary analyses are presented in Figure 7 and Figure S4. Interestingly, even with a 10-fold higher MOI used for the WT infection of donor Vero-ACE2 cells relative to other variants, we observed comparable rates of cell-to-cell transmission between WT, D614G, B.1.1.7, and B.1.351 (Figure 7A, upper panel; Figure 7B and Figure S4A). Note that the relative rate of cell-to-cell transmission shown in Figure 7B was obtained by dividing the percentage of SARS-CoV-2-positive Vero-mTomato-Red cells (*Q2* in Figure 7A, upper panel) by the percentage of total SARS-CoV-2-positive cells (Q2 plus *Q*3 in Figure 7A, upper panel). We noted that the rate of B.1.351 spreading infection in Vero-ACE2 and Vero-mTomato-Red cells (Q2 plus Q3 in **Figure A**, upper panel) was the highest, followed by B.1.1.7 > D614G > WT (Figure 7C). Consistent with the more efficient replication of B.1.351 in donor Vero-ACE2 cells over the entire 26 hr infection period (Q3 in Figure 7A, upper panel), we found a significantly higher cell-free infectivity for B.1.351 produced during the initial 20-hr infection relative to WT, D614G and B.1.1.7 (Figure 7D, see “no sera”). Overall, these results revealed a strongly enhanced replication of B.1.351 relative to B.1.1.7, D614G and WT, yet a comparable efficiency of cell-to-cell transmission between WT, D614G and VOCs.

**Figure 7.**
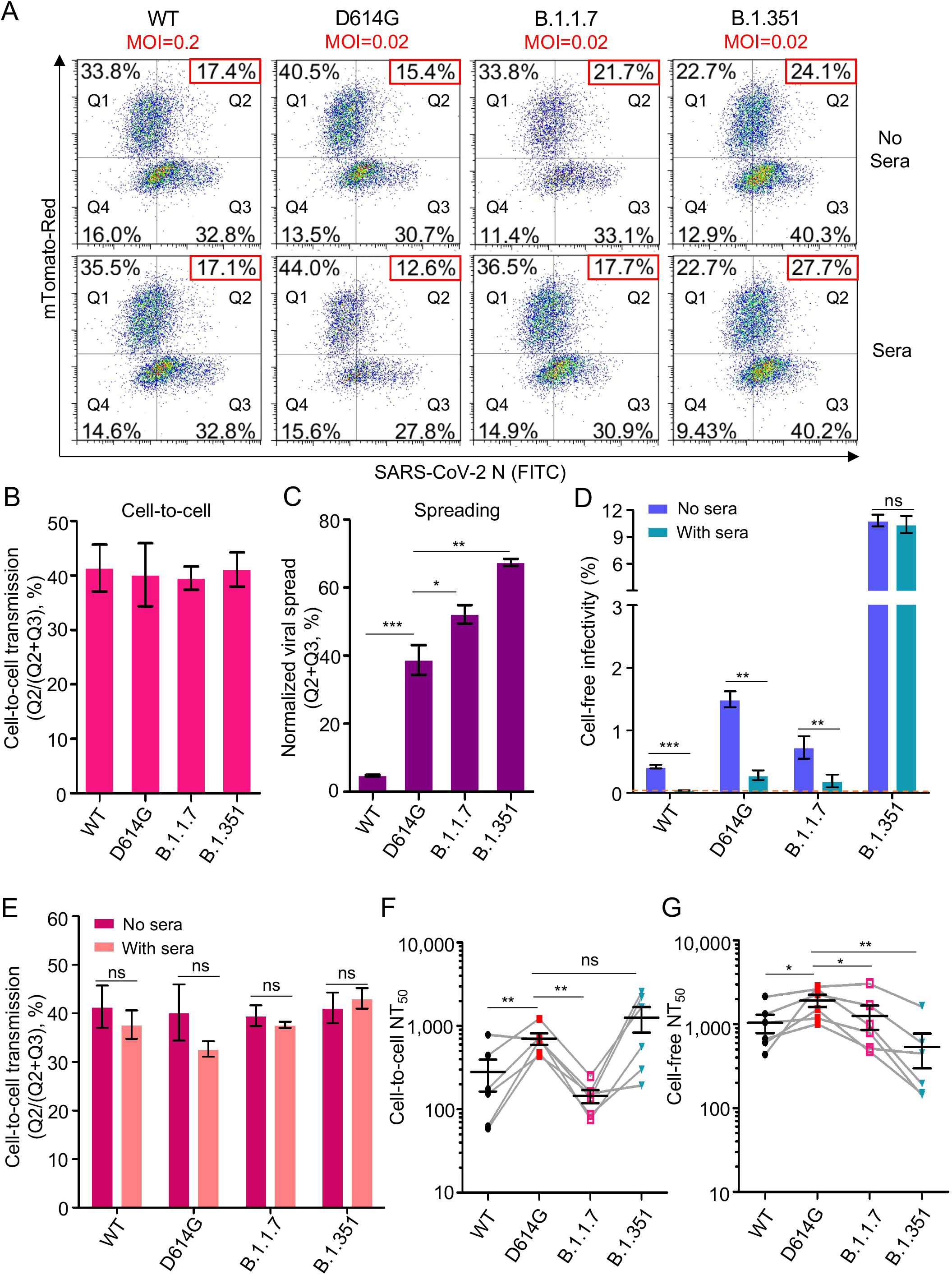
Cell-to-cell transmission of SARS-CoV-2 VOCs and sensitivity to neutralization by vaccinee sera. **(A-E)** The cell-to-cell transmission capability of authentic SARS-CoV-2 WT, D614G, B.1.1.7 and B.1.351 in the presence or absence of vaccinee sera. Donor Vero-ACE2 cells were infected with WT SARS-CoV-2 (MOI=0.2), D614G (MOI=0.02), B.1.1.7 (MOI=0.02), and B.1.351 (MOI=0.02) for 20 hr, followed by coculturing with target Vero-mTomato (Red) cells in the presence or absence of pooled mRNA vaccinee sera (3 Moderna and 3 Pfizer) for 6 hr. Cells were fixed and stained with anti-SARS-CoV-2 N protein, and analyzed by flow cytometry. Representative flow cytometric analyses of infected cells were shown in (**A**), with the newly infected target Vero-mTomato (Red) cells (Q2) as indicative of cell-to-cell transmission. The relative cell-to-cell transmission efficiency was calculated by dividing the rate of Vero-mTomato-Red positive cells (Q2) by the rate of total infected donor and target cells (Q2+Q3) (**B,** n=3). The MOI-normalized total viral spread in both donor and target cells (Q2+Q3) was shown in **(C)** (n=3). The supernatant from the initial 20 hr infection of donor cells was used to infect target Vero-mTomato-Red cells for 6 hr as the measurement of cell-free viral infectivity, either in the presence or absence of the pooled vaccinee sera, and infected cells were analyzed by flow cytometry **(D)** (n=3). The pooled vaccinee sera were also added to the cocultured Vero-ACE2 and Vero-mTomato-Red cells as described in (**A**) to determine their effect on cell-to-cell transmission **(E)**. **(F and G)** The calculated NT_50_ values of vaccine sera against cell-to-cell transmission and cell-free infection of lentiviral pseudotypes bearing individual spike of VOCs. Experimental procedures were the same as described in Figures 6D and 6E, except that all comparisons were made relative to the D614G variant (n=6).

We also assessed the sensitivity of cell-to-cell transmission and cell-free infection to neutralization by Moderna and Pfizer vaccinee sera. With a relatively low dose of pooled sera being applied, we observed that the cell-to-cell transmission of WT, D614G, B.1.1.7 and B.1.351 was virtually resistant to neutralizing antibodies induced by these mRNA vaccinees for all viruses, whereas the cell-free infection of WT, D614G and B.1.1.7 was strongly inhibited, with B.1.351 being resistant (Figure 7A, lower panels; Figure 7D **and** Figure 7E; Figure S4B **and** Figure S4C). By using HIV-inGluc pseudotyped viruses with serially diluted serum samples from Moderna and Pfizer vaccinees, we were able to obtain and compare the NT_50_ values of each virus in cell-to-cell transmission vs. cell-free infection. We found that, overall, mRNA vaccinee sera neutralized cell-to-cell transmission approximately 3-fold less efficiently than cell-free infection, with the notable exception of B.1.351, which showed similar extents of inhibition for cell-to-cell and cell-free infections (Figure 7F **and** Figure 7G). Intriguingly, we found that the cell-to-cell transmission of B.1.1.7 was more resistant to neutralization by vaccine sera, with ∼4.9-fold lower NT_50_ than D614G (p<0.01) and ∼8.7-fold lower than B.1.351 (p<0.05) (Figure 7F **and** Figure 7G). In contrast, the cell-free infection of B.1.351 was more resistant to neutralization than D614G and B.1.1.7, with 3.6-fold (p<0.01) and ∼2.4-fold (p<0.01) lower NT_50_, respectively (Figure 7F **and** Figure 7G), which was consistent with recent studies (Planas et al., 2021; Wang et al., 2021). In aggregate, these results confirmed that cell-to-cell transmission of both authentic and pseudotyped SARS-CoV-2 VOCs is more refractory to inhibition by neutralizing antibodies induced by mRNA vaccines as compared to cell-free infection, and more importantly, showed that the cell-to-cell transmission of B.1.1.7 and the cell-free infection of B.1.351, are most resistant to the antibody neutralization. The differential sensitivity of B.1.1.7 and B.1.351 to neutralization by vaccinee sera in cell-to-cell transmission vs cell-free infection likely has important implications for understanding the spread of these variants and their pathogenesis in patients (see Discussion).

## Discussion

Accumulating evidence indicates that viruses, including the highly pathogenic HIV, HCV, and EBOV, etc., can efficiently spread through cell-to-cell transmission (Cifuentes-Munoz et al., 2018; Dale et al., 2013; Miao et al., 2016; Sattentau, 2008; Wang et al., 2017; Xiao et al., 2014; Zhong et al., 2013a). Importantly, cell-to-cell transmission is more efficient than cell-free infection (Zhong et al., 2013a), and roles for this mode of transmission have been demonstrated in vivo for HIV and other viruses (Agosto et al., 2014; Dale et al., 2013; Xiao et al., 2014; Zhong et al., 2013a). Notably, many plant viruses are known to use cell-to-cell transmission to spread from epidermal cells and move sequentially into mesophyll, bundle sheath, and phloem parenchyma and companion cells (Carrington et al., 1996; Hipper et al., 2013). For coronaviruses, very little is currently known about their mode of spread between cells or its efficiency compared to cell-free infection. This question is critical, given the robust replication of SARS-CoV-2 in human lung and other tissues, as well as the rapid spread of SARS-CoV-2, including some variants of concern, in the human population, leading to the global pandemic (Chu et al., 2020; Grubaugh et al., 2021; Planas et al., 2021; Walensky et al., 2021; Wang et al., 2021). In this work, we addressed this question using lentiviral pseudotypes and replication-competent rVSV expressing the spike of SARS-CoV-2 or SARS-CoV. We discovered that SARS-CoV-2 spike is more efficient in mediating cell-to-cell transmission than SARS-CoV spike, yet the spike of SARS-CoV is more capable of mediating cell-free infection. To our knowledge, this is the first direct comparison of cell-to-cell transmission vs. cell-free infection between SARS-CoV-2 and SARS-CoV in cultured cells, and the results provide important insights into two distinct modes of infection and the host-viral factors that regulate these processes.

Why is SARS-CoV-2 spike more efficient than SARS-CoV spike for mediating cell-to-cell transmission in cultured cells? We provide evidence that this is in part related to the higher cell-cell fusion activity of SARS-CoV-2 spike compared to SARS-CoV (Figure 2). However, we also recognized that extensive cell-cell fusion by SARS-CoV-2 spike can lead to giant syncytia formation and cell death, which in turn reduces cell-to-cell transmission. Therefore, fine control of the spike-induced cell-cell fusion is important for efficient cell-to-cell transmission and, therefore, the spreading infection of SARS-CoV-2. Further evidence supporting a role of cell-cell fusion in transmission of SARS-CoV-2 came from the application of a membrane fusion inhibitor EK1, which significantly attenuated cell-to-cell transmission. Interestingly, although ACE2 enhances cell-to-cell transmission of SARS-CoV-2 and SARS-CoV, we found that it is not absolutely required. This observation is supported further by data from H520 cells and human PBMCs, which express a minimal level of ACE2 if any, yet exhibited obvious cell-to-cell transmission (Figure 4). Cell-free infection of SARS-CoV-2 was not detected in H520 cells and PBMCs, further supporting these conclusions. The molecular mechanism underlying cell-to-cell transmission of SARS-CoV-2, including the possible roles of cellular cofactors and virological synapses, shall be investigated in future studies.

A surprising result to emerge from our studies was that, despite the critical role of cell-cell contact and plasma membrane-mediated fusion, endosomal entry pathways were also involved in cell-to-cell transmission of SARS-CoV-2 and SARS-CoV (Figure 5). This is evidenced by the inhibitory effect of drugs that specifically target the endosomal entry pathway of these viruses, including the CatL inhibitor III, which blocks cleavage of the viral glycoprotein, as well as BafA1, which neutralizes endosomal pH. These results are reminiscent of previous studies from HIV and EBOV, where endocytosis and/or protease cleavage are required for cell-to-cell transmission of these enveloped viruses (Dale et al., 2011; Markosyan et al., 2016; Miao et al., 2016; Titanji et al., 2013; Wang et al., 2017). Interestingly, we find that these inhibitors appear to be less potent for decreasing cell-to-cell transmission as compared to cell-free infection, and moreover, their effects on SARS-CoV-2 are less than their effects on SARS-CoV. These observations collectively suggest a less dominant role for the endosomal entry pathway in cell-to-cell transmission of SARS-CoV-2. High-resolution live microscopic imaging would be useful to dissect the exact role of endosomal vs. plasma entry pathway in the cell-to-cell transmission of SARS-CoV-2.

Cell-to-cell transmission is considered to be an effective means by which viruses evade host immunity, especially antibody-mediated responses. We compared the sensitivity of cell-to-cell transmission vs. cell-free infection of SARS-CoV-2 to treatments by neutralizing monoclonal antibodies and COVID-19 convalescent plasma - both of which have been approved by the FDA for emergency use. We found that while cell-free infection of SARS-CoV-2 was almost completely blocked by these treatments, cell-to-cell transmission of SARS-CoV-2 was, to a large extent, refractory (Figures 6 and 7). While not statistically significant, some of the COVID-19 sera (2 out of 5) even enhanced cell-to-cell transmission of SARS-CoV-2 (Figure 6D), although the underlying mechanisms are currently not known. Interestingly, despite significant increases in cell-free infectivity, the South Africa variant B.1.351, the UK variant B.1.1.7, as well as the D614G variant, exhibited similar efficiencies of cell-to-cell transmission compared with the WT (Figure 7). Moreover, although B.1.351 is more resistant to vaccinee sera in cell-free infection, consistent with some recent reports (Planas et al., 2021; Wang et al., 2021), B.1.1.7 seems more resistant to the vaccinee sera for the cell-to-cell transmission route (Figure 7), may explain that B.1.1.7 has longer duration of acute infection than others (Kissler et al., 2021). The mechanism underlying these observations is currently unclear, but may have implications for understanding the rapid spread of VOCs in human population as well as their increased pathogenesis. The cell-free route is directly linked to the ability of viruses to infect target cells and result in spreading among humans through person-to-person contact. In contrast, cell-to-cell transmission has dominant roles in viral pathogenesis and disease progression (Mothes et al., 2010). Thus, our results on the resistance of B.1.1.7 and B.1.351 to vaccinee sera-mediated inhibition of cell-to-cell transmission and cell-free infection may provide molecular and virological underpinnings for the prolonged viral replication and rapid spread of these two variants in the world population (Alpert et al., 2021; Funk et al., 2021; Planas et al., 2021; Wang et al., 2021).

### Limitations of the Study

While in this work, we obtained evidence that SARS-CoV-2 spike more efficiently mediates cell-to-cell transmission than the SARS-CoV spike, a direct comparison using authentic viruses of both, especially in primary human lung and airway epithelial cells, is needed. As an initial step toward this goal, we have attempted to apply rVSV-GFP-SARS-CoV-2 and rVSV-GFP-SARS-CoV to human primary bronchial and nasal epithelial cell cultures, but the efficiency of spread for both viruses was too low to draw any conclusion. Although in this work, we examined roles of ACE2 and endosomal proteases in cell-to-cell transmission vs. cell-free infection, how other host cofactors, including TMPRSS2, modulate this process will need to be investigated. Ultimately, we must determine the role, if any, of cell-to-cell transmission of SARS-CoV-2 in disease progression and pathogenesis in COVID-19 patients.

## ACKNOWLEDGEMENTS

We thank Gerard Lozanski, Richard Gumina, Eric Freed, David Derse, Marc Johnson, Fang Li, and Ali Ellebedy for provision of sera samples, plasmids, and cells. We also thank the NIH AIDS Reagent Program and BEI Resources for supplying important reagents that made this work possible. This work was supported by a fund provided by an anonymous private donor to The Ohio State University and NIH grant U54CA260582; additional support of SLL’s lab includes NIH grants R01 AI112381 and R01 AI150473. The content is solely the responsibility of the authors and does not necessarily represent the official views of the NIH. LJS was partially supported by NIH grant NICHD R01 HD095881.

## Author contributions

SLL conceived and led the project. CZ performed majority of the presented experiments. JPE designed the construction of variants of concern and performed part of the neutralization assay. TK performed rVSV-GFP spread experiments in human airway epithelial cells. YMZ produced rVSV viruses and sequenced the spike gene. SPJW contributed to the rVSV stock. CZ, JPE, TK, YMZ, GO, SPJW, LS, MEP and SLL all contributed to data analyses and discussion. CZ and SLL wrote the paper, which was edited and approved by all coauthors.

## STAR★METHODS

### LEAD CONTACT AND MATERIALS AVAILABILITY

Further information and requests for resources and reagents should be directed to and will be fulfilled by the Lead Contact, Shan-Lu Liu.

### EXPERIMENTAL MODEL AND SUBJECT DETAILS

#### Cell culture

293T (ATCC CRL-11268, RRID: CVCL_1926), Vero-E6 (ATCC CRL-1586, RRID: CVCL_0574) and Vero-ACE2 (Vero-E6 expressing high endogenous ACE2, BEI, NR-53726) cells were grown in Dulbecco’s modified Eagle’s medium (DMEM) supplemented with 1% penicillin/streptomycin and 10% (vol/vol) fetal bovine serum (Thermo Fisher Scientific). Caco-2 (ATCC HTB-37, RRID: CVCL_0025) cells were grown in Dulbecco’s modified Eagle’s medium (DMEM) supplemented with 1% penicillin/streptomycin and 20% (vol/vol) FBS. Calu-3 cells (ATCC HTB-55, RRID: CVCL_0609) were grown in Eagle’s Minimum Essential Medium (EMEM) supplemented with 1% penicillin/streptomycin and 10% (vol/vol) FBS. Human peripheral blood mononuclear cells (PBMCs) were gifts of Eric O. Freed (National Cancer Institute, Frederick, Maryland, USA) and maintained in Roswell Park Memorial Institute (RPMI) 1640 Medium containing 10% (vol/vol) FBS. NCI-H520 (ATCC HTB-182, RRID: CVCL_1566) cells were grown in RPMI 1640 Medium supplemented with 1% penicillin/streptomycin and 10% (vol/vol) fetal bovine serum. The 293T/ACE2 cell line was obtained from BEI (NR-52511). Vero-E6 cells stably expressing red tomato were generated by transduction of a lentiviral vector expressing the tomato gene, followed by hygromycin B selection (200 µg/mL) for 6 days. All cell lines utilized were maintained at 37°C, 5% CO2.

#### Virus

rVSV-GFP-SARS-CoV and rVSV-GFP-SARS-CoV-2 (obtained from Sean Whelan’s lab at the Washington University School of Medicine in St. Louis, Missouri, USA) were amplified in Vero-E6 cells and maintained under a humidified atmosphere of 5% CO_2_ at 34°C in Dulbecco’s modified Eagle’s medium (DMEM) supplemented with 10% FBS. The spike sequence in the original stock and each passage was confirmed by DNA sequencing. Authentic SARS-CoV-2 WT (USA-WA1/2020, NR-52281; kindly prepared by Jacob Yount of The Ohio State University, Columbus, Ohio, USA), D614G (B.1.5, NR-53944), B.1.1.7 (501Y.V1, NR-54000) and B.1.351 (501Y.V2, NR-54009) were all obtained from BEI.

### METHOD DETAILS

#### Constructs, antibodies and reagents

HIV-1 NL4.3-inGluc was a gift of Marc Johnson at the University of Missouri (Columbia, Missouri, USA). Plasmids pcDNA3.1-SARS-CoV-S-C9 and pcDNA3.1-SARS-CoV2-S-C9 encoding the full-length spike were obtained from Fang Li at the University of Minnesota (St. Paul, Minnesota, USA). A construct for ACE2 transient expression, pHAGE2-ACE2, was obtained from BEI resources (NR-52512). A lentiviral vector encoding red tomato was from Marc Johnson (University of Missouri, Columbia, USA). The codon-optimized D614G, B.1.351 and B.1.1.7 SARS-CoV-2 S constructs were synthesized by GenScript and subsequently cloned into a pcDNA3.1 vector by restriction enzyme cloning with Kpn I and BamH I. Primary antibodies used for western blotting and flow cytometry were anti-coronavirus spike (Sino Biological, 40150-T62), anti-SARS-CoV-2 Nucleocapsid (Sino Biological, 40143-MM08), anti-hACE2 (R&D, AF933) and anti-β-actin (Sigma, A1978). Secondary antibodies used for western blotting included anti-Mouse IgG-Peroxidase (Sigma, A5278), anti-Rabbit IgG-Peroxidase (Sigma, A9169) and anti-Goat IgG-Peroxidase (Sigma, A8919). Secondary antibodies used for flow cytometry included anti-Rabbit IgG–FITC (Sigma, F9887), anti-Mouse IgG-FITC (Sigma, F0257), anti-Goat IgG-FITC (Sigma, F7367). The monoclonal Ab 2B04 was a gift of Ali Ellebedy (Washington University in St. Louis).

Inhibitors in this study included Methyl cellulose (Sigma, M0512), Cathepsin L Inhibitor III (Sigma, 219427), CA-074 Me (Sigma, 205531), EST/E-64D (Sigma, 330005), Bafilomycin A1 (Sigma, B1793) and Leupeptin (Sigma, L2884). EK1 peptide was synthesized by Alpha Diagnostic International (San Antonio, Texas).

Patient serum samples were collected from hospitalized COVID-19 patients under The Ohio State University IRB protocol #2020H0228 as described (Zeng et al., 2020). Vaccinee serum samples were collected from health care workers following 3-4 weeks of the second dose of Moderna and Pfizer SARS-CoV-2 mRNA vaccination under an amended IRB protocol #2020H0228.

#### Cell-to-cell transmission

In the lentiviral vector system, the expression of anti-sense reporter gene Gluc is interrupted by an intron oriented in the sense direction of the HIV-1 genome so that Gluc production will only occur in infected target cells and not virus producer cells (Zeng et al., 2020). By coculturing the virus producer and target cells, cell-to-cell transmission was determined by measuring the Gluc activity of the cocultured media between donor cells (such as 293T) producing lentiviral pseudotypes and target cells (such as 293T/ACE2). Specifically, 293T cells were seeded in 6-well plates and transfected with 1.4 µg NL4.3-inGluc and 0.7 µg of plasmids encoding SARS-CoV or SARS-CoV-2 spike. The next day, transfected 293T donor cells were digested with PBS/5 mM EDTA and thoroughly washed with PBS to remove EDTA, followed by coculturing with target cells (293T/ACE2, Caco-2, Calu-3, NCI-H520 or PBMCs) at 1:1 ratio in 24-well plates for 24∼72 hr. Inhibitors or sera were added as needed. Supernatants were collected and measured for the Gluc activity.

For authentic SARS-CoV-2 WT and VOCs, the donor Vero-ACE2 cells were infected with an MOI of 0.2 (WT) or 0.02 (VOCs) for 20 hr, followed by coculturing with the same number of Vero-mTomato-Red cells for an additional 6 hr, in the presence or absence of vaccinee sera. Cells were then fixed with 3.7% formaldehyde for 1 hr, followed by three times of wash with PBS before being taken out of the BSL3 lab. The fixed cells were incubated with anti-SARS-CoV-2 Nucleocapsid and anti-Mouse-FITC, and subjected to flow cytometry analysis.

#### Cell-free infection

Cell-free infection was performed along with cell-to-cell transmission in this work. Briefly, an equal number of transfected donor cells were seeded in new 24-well plates and maintained for the same period of time as in cell-to-cell transmission (normally 48-72 hr). The total volumes of supernatants were collected and used to infect target cells, which were seeded with the presence of the same amount of untransfected 293T cells; this would ensure that the total numbers of cells and density used for cell-to-cell and cell-free infection assays were comparable. For the *transwell* setting, the transfected donor cells were seeded onto the insert while target cells, which again were mixed with same amount of untransfected 293T cells, were on the bottom; this would avoid the contact between donor and target cells yet the virus can spread through the filter. Supernatants were collected at the same time points as cell-to-cell transmission and measured for Gluc activity.

#### Cell-cell fusion

For fluorescence-based cell-cell fusion, 293T cells were transfected with plasmid encoding GFP and spikes. Following 24 hrs transfection, donor 293T cells were cocultured with target cells. Micrographs of cocultured cells were taken after 2∼24 hrs coculture. For tet-off-based assay, 293T cells were transfected with plasmids encoding tet-off or SARS-CoV or SARS-CoV-2 spike and were cocultured with target 293FT-mCAT-Gluc cells (stably expressing tetracycline-responsive element (TRE)-driven *Gaussia* luciferase), which were transfected with a plasmid expressing ACE2; Gluc activity was measured from the supernatant of cocultured cells harvested at 24 and 48 hr, respectively.

#### Plaque assay

The replication-competent rVSV-GFP-SARS-CoV and rVSV-GFP-SARS-CoV-2 viruses were used to infect confluent Vero-E6 cells (MOI=0.01) for 1 h at 37°C. The uninfected virus was then removed from cells and replaced with 1% methylcellulose in DMEM/5% FBS and incubated for 72 hr at 37°C. Cells were fixed with 3.7% paraformaldehyde in PBS and stained with 1% crystal violet (Sigma, C0775) in 10% ethanol for visualization of plaques.

#### Flow cytometry

For analysis of spike and ACE2 expression on the cell surface, transfected 293T cells were washed with PBS, detached with PBS/5mM EDTA for 10 min, washed twice with cold PBS/2% FBS, and incubated with anti-coronavirus Spike/Nucleocapsid or anti-hACE2 antibody for 1 hr. After three washes with cold PBS/2% FBS, cells were incubated with FITC-conjugated anti-rabbit IgG/anti-mouse IgG or anti-goat IgG (1:200) secondary antibodies for 1 hr. Cells were washed three times with cold PBS /2% FBS and fixed with 3.7% formaldehyde for 10 min and analyzed by flow cytometry. For analysis of rVSV-GFP-SARS-CoV and rVSV-GFP-SARS-CoV-2 infection, infected Vero E6 cells were washed with PBS and digested with 0.05% trypsin, followed by fixation with 3.7% formaldehyde for 10 min and analyzed by flow cytometry.

#### Western blotting

Western blotting was performed as previously described (Li et al., 2019; Zeng et al., 2020). In brief, HEK293T cells were collected and lysed in RIPA buffer (50 mM Tris, pH 7.5, 150 mM NaCl, 1 mM EDTA, 1% Nonidet P-40, 0.1% SDS, protease inhibitor cocktail) for 40 min on ice, followed by centrifugation for 10 min, 12,000 x g at 4°C, Cell lysate then boiled at 100 ℃ for 10 min with 1XSDS loading buffer containing 2-Mercaptoethanol. Samples were run on 10% SDS-PAGE gels, transferred to PVDF membranes, and probed with primary antibodies and secondary antibodies, analyzed by Amersham Imager 600 (Thermofisher).

#### Neutralization assays

Cell-free virus neutralization assays were performed by incubating free virus with serial diluted Moderna and Pfizer vaccinee sera, followed by infecting 293T/ACE2 target cells and measuring the luciferase activity (Zeng et al., 2020) at 48 and 72 hr. Cell-to-cell virus neutralization assays were performed by incubating serial diluted sera with viral producer cells (transfected 293T) and target cells (293T/ACE2) in the coculture system, and supernatants were collected at 48 and 72 hr to measure the luciferase activity. In both cases, NT_50_ was defined as the sera dilution fold at which the relative light units were reduced by 50% compared with the control wells (no sera); the NT_50_ values were calculated using nonlinear regression in GraphPad Prism.

### QUANTIFICATION AND STATISTICAL ANALYSIS

#### Statistical Analysis

Data were analyzed as mean with Standard Error of Mean (SEM). All experiments were performed at least three independent replications, and the number of biological replicates for each data set is given by “n” and is provided in the respective figure legend. Statistical analyses were performed using GraphPad Prism 5.0 as follows: One-way Analysis of Variance (ANOVA) with Bonferroni’s post-tests was used to compute statistical significance between multiple groups for multiple comparison or t-test was used for two groups for single comparison. A p value of less than 0.05 was considered significant and indicated by an asterisk (*, p<0.05).

## Supplemental Figures

**Figure S1.**
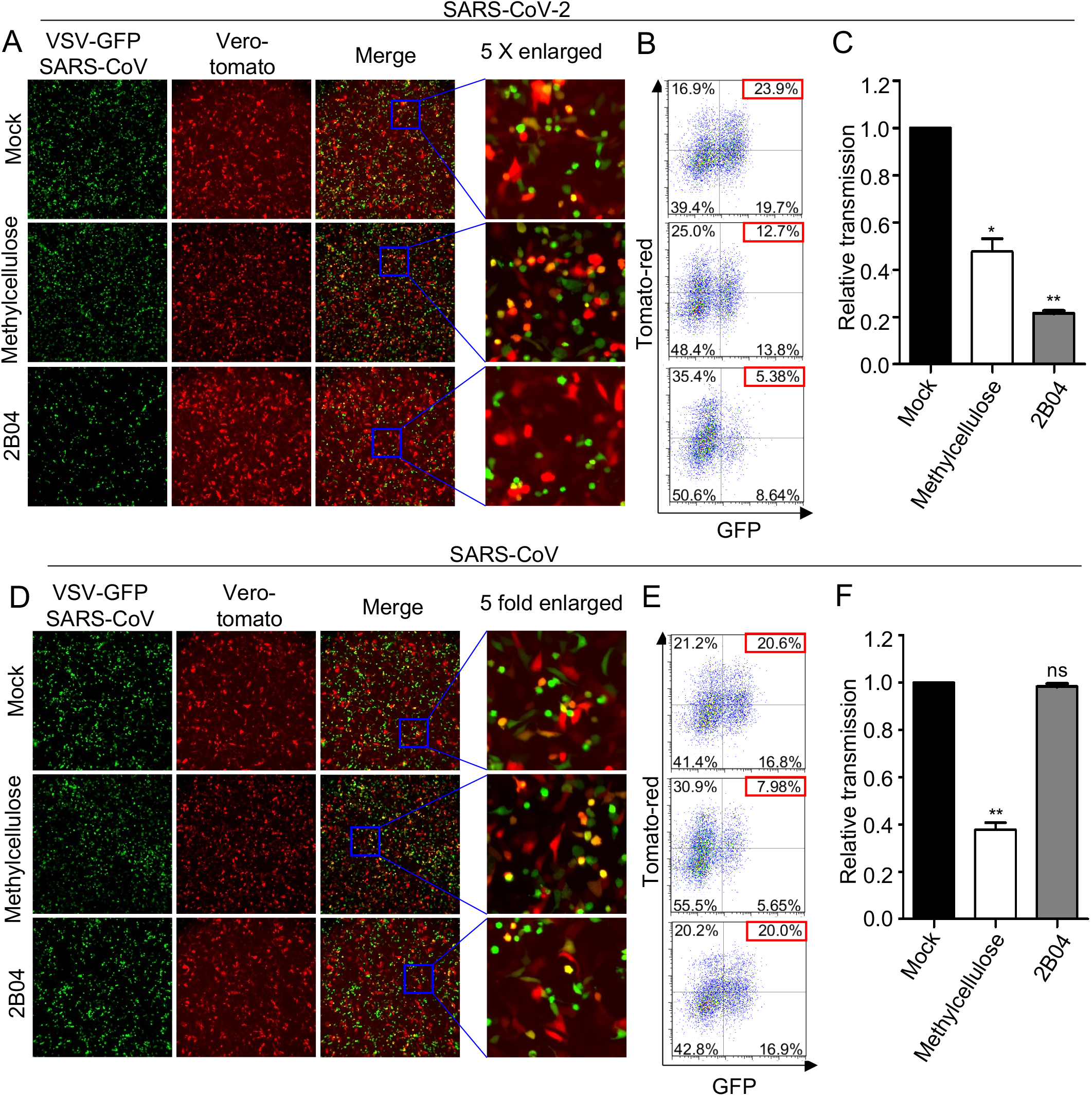
Effects of methylcellulose and monoclonal antibody 2B04 on rVSV-GFP transmission in Vero-E6 cells. Vero-E6 cells were infected with appropriate MOIs of either VSV-GFP-SARS-CoV or VSV-GFP-SARS-CoV-2. After 16 h post-infection, the infected Vero-E6 cells were cocultured with Vero-mTomato-Red cells at 1:1 ratio, in the presence or absence of 2 µg/mL 2B04 or 1% methylcellulose. Micrographs of cocultured cells were taken after 18 h coculture **(A and D)**, with dual fluorescence positive cells indicated by arrows. The GFP signals in Tomato-positive cells were analyzed by flow cytometry **(B and E**, Q2), indicative of virus transmission from Vero-E6 to Vero-mTomato-Red cells. Results from 3 independent experiment (n=3) were summarized and plotted as relative infection rates by setting the values of mock infection control to 1.0 (**C and F**).

**Figure S2.**
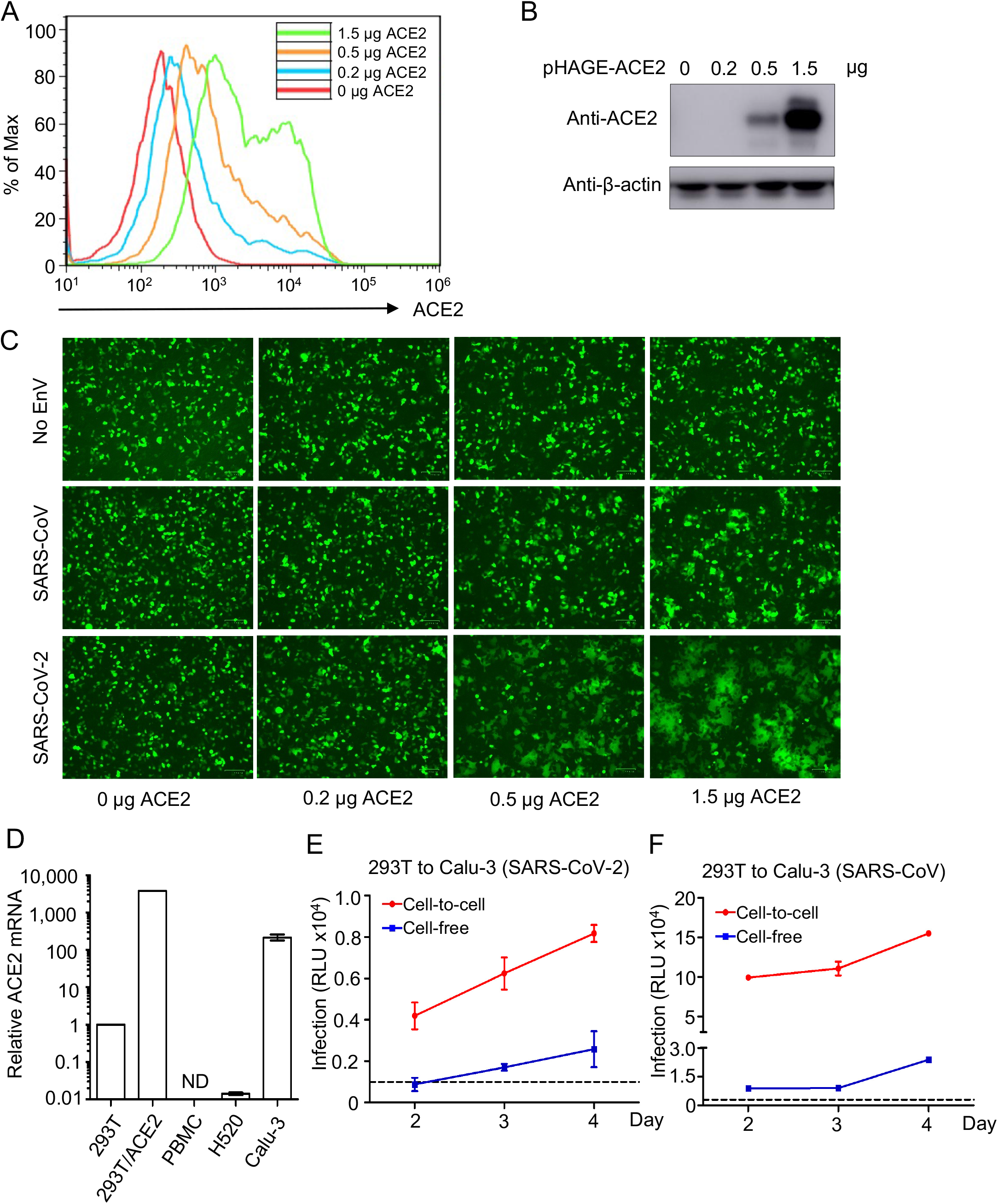
Role of ACE2 in cell-to-cell transmission. **(A and B)** The expression level of ACE2 in target cells was analyzed by flow cytometry **(A)** and western blotting **(B)** using a specific antibody against ACE2; results were one representative of three independent experiments. **(C)** Representative images of cell-cell fusion induced by SARS-CoV-2 and SARS-CoV spike at indicated doses of ACE2. **(D)** The expression level of ACE2 in different cell lines and human PBMCs. qPCR was performed to quantify the ACE2 mRNA expression and relative expression was plotted by setting the value of 293T cells to 1.0. ND: not detected. **(E and F)** Cell-to-cell transmission in Calu-3 cells. Experiments were performed as described in Figures 1 and 4, except that Calu-3 cells were used as target cells, which were cocultured with viral producer 293T cells (n=3).

**Figure S3.**
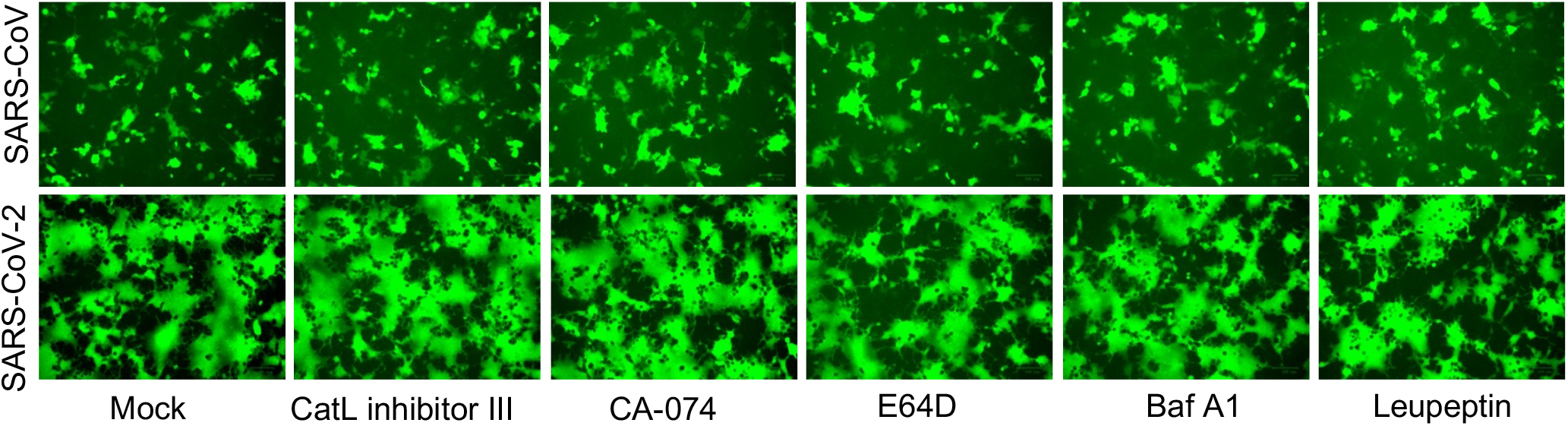
Effect of endosomal entry inhibitors on cell-cell fusion induced by SARS-CoV-2 spike. Experiments were carried out as described in Figures 3 and 5, with indicated inhibitors included in the cell coculture: 5 µM Cat L inhibitor III, 5 µM CA-074, 30 µM E-64D, 50 nM BafA1, and 50 µM leupeptin.

**Figure S4.**
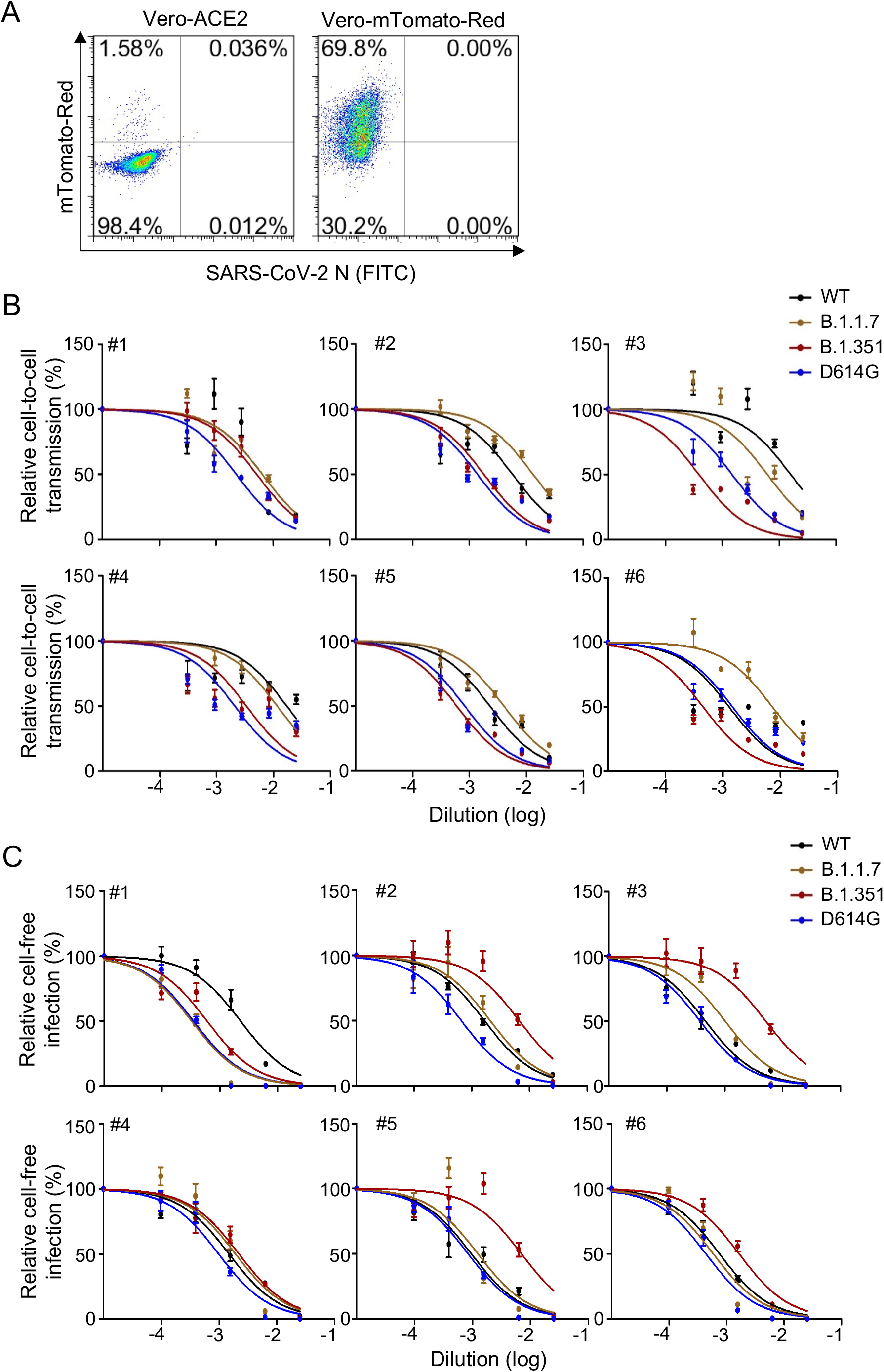
Neutralization curves of vaccinee sera against the cell-to-cell and cell-free infection of VOCs B1.1.7 and B.1.351 relative to D614G and WT. **(A)** Flow cytometric gating control in analysis of data presented in Figure 7A using uninfected Vero-ACE2 and Vero-mTomato-Red cells. **(B and C)** Six vaccinee sera samples, 3 from Moderna and 3 from Pfizer, were chosen for the neutralization assay in the context of cell-to-cell transmission or cell-free infection. The y axis indicates the relative viral infectivity by setting the viral infectivity without serum to 100%; the x axis indicates dilution fold of serum samples (n=6).

## Highlights

1. SARS-CoV-2 spike efficiently mediates cell-to-cell transmission
2. Cell-cell fusion promotes cell-to-cell transmission of SARS-CoV-2
3. ACE2 enhances but is not essential for cell-to-cell transmission
4. Cell-to-cell transmission of SARS-CoV-2 is resistant to Ab neutralization

## In Brief

The spike protein of SARS-CoV-2 mediates cell-to-cell transmission that is promoted by cell-cell fusion. ACE2 enhances cell-to-cell transmission but is not essential. Cell-to-cell transmission of SARS-CoV-2 is refractory to antibody neutralization.

## Graphical Abstract

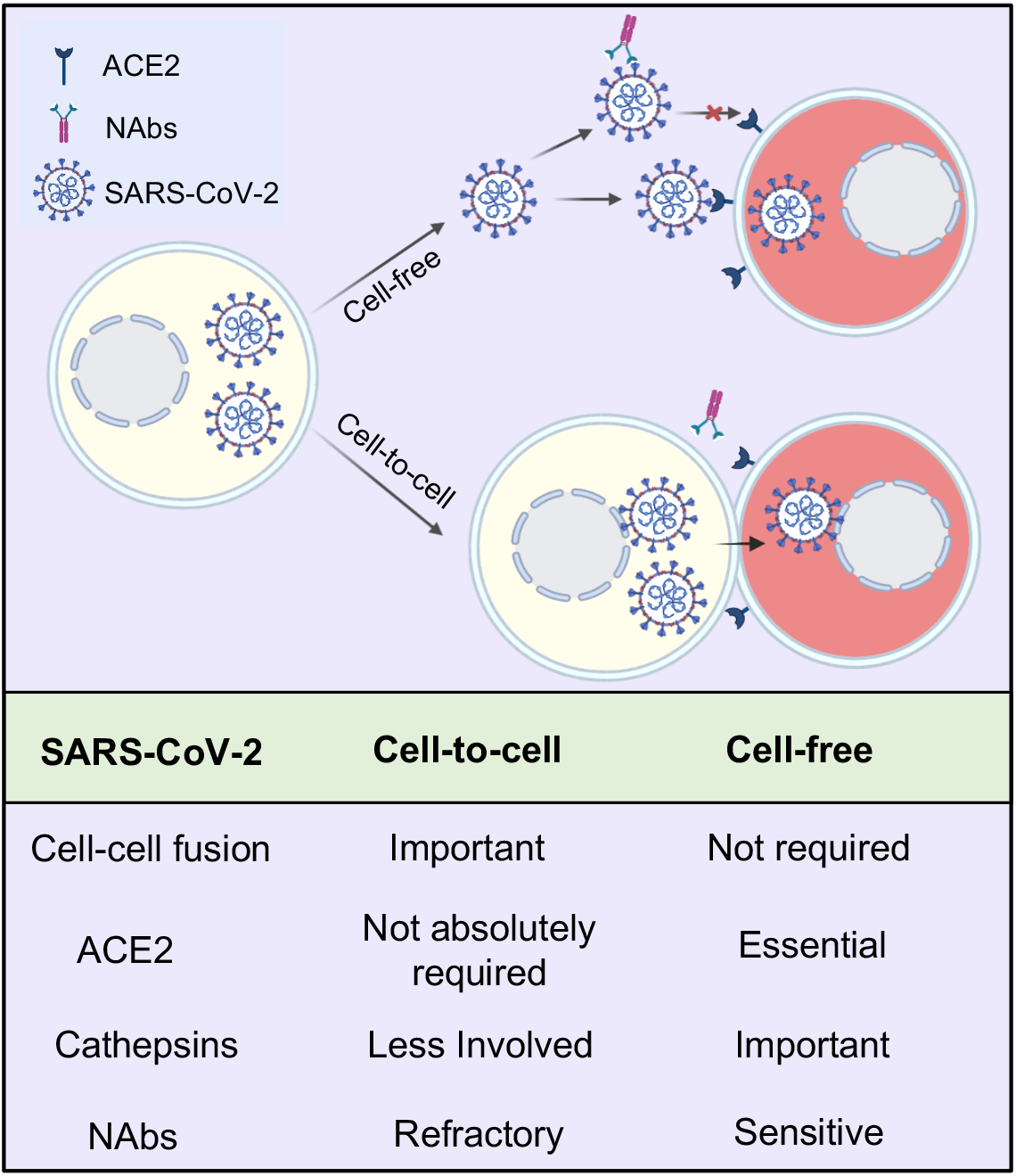

